# Host identity and symbiotic association affects the genetic and functional diversity of the clownfish-hosting sea anemone microbiome

**DOI:** 10.1101/745760

**Authors:** Benjamin M. Titus, Robert Laroche, Estefanía Rodríguez, Herman Wirshing, Christopher P. Meyer

## Abstract

All eukaryotic life engages in symbioses with a diverse community of bacteria that are essential for performing basic life functions. In many cases, eukaryotic organisms form additional symbioses with other macroscopic eukaryotes. The tightly-linked physical interactions that characterize many macroscopic symbioses creates opportunities for microbial transfer, which likely affects the diversity and function of individual microbiomes, and may ultimately lead to microbiome convergence between distantly related taxa. Here, we sequence the microbiomes of five species of clownfish-hosting sea anemones that co-occur on coral reefs in the Maldives. We test the importance of evolutionary history, clownfish symbiont association, and habitat on the genetic and predicted functional diversity of the microbiome, and explore signals of microbiome convergence in anemone taxa that have evolved symbioses with clownfishes independently. Our data indicate that host identity shapes the majority of the genetic diversity of the clownfish-hosting sea anemone microbiome, but predicted functional microbial diversity analyses demonstrate a convergence among host anemone microbiomes, which reflect increased functional diversity over individuals that do not host clownfishes. Further, we identify up-regulated microbial functions in host anemones that are likely affected by clownfish presence. Taken together our study reveals an even deeper metabolic coupling between clownfishes and their host anemones, and what could be a previously unknown mutualistic benefit to anemones that are symbiotic with clownfishes.

## Introduction

The importance of symbiosis is underscored by its ubiquity-virtually all of life engages in complex multi-level symbioses that critically impact the formation and distribution of biodiversity around the globe [1-3]. Minimally, all multicellular life engages in symbioses with prokaryotic microbiota that are essential for survival, critically impact individual health, development, and nutrient acquisition, and which serve as a primary interface between individuals and their environment [e.g. 4-12]. The composition and diversity of an individual’s microbial community (i.e. microbiome) has been shown to be influenced by a variety of factors, but are generally discussed as a combination of evolutionary history and ecology [e.g. 13-17].

In addition to microbial symbioses, many multicellular eukaryotes engage in symbioses with other multicellular organisms. Central features of these interactions are the physical linkages between constituent partners which provide opportunity for microbial transfer, adding a layer of complexity to the factors that affect the diversity and function of individual microbiomes. Many macroscopic symbioses involve diverse partners, occur across distinct habitats, and have evolved independently multiple times across the tree of life [18-20]. Consequently, macroscopic symbioses provide an important framework for exploring the processes that may lead distantly related taxa to converge on microbiomes with similar genetic and functional profiles.

Among the immense symbiotic diversity on the planet, the clownfish-sea anemone mutualism stands out as an iconic example, and holds characteristics that make it a useful system for understanding the processes that affect microbiome diversity and function within macroscopic symbioses. A classic example of mutualism, the 30 species of clownfishes (or anemonefishes) have adaptively radiated from a common ancestor to live with 10 species of sea anemones on coral reefs of the Indian and Pacific Oceans [21-23]. Within sea anemones (Order Actiniaria), symbiosis with clownfishes has evolved independently at least three times [24]. The clownfish-sea anemone symbiosis is found across a range of coral reef habitats and involves many combinations of sea anemone-clownfish associations that are co-distributed [21, 23-27]. Clownfishes are considered obligate symbionts of sea anemones, have evolved a range of host specificities, and are never found solitarily [21, 27]. Anemones in contrast, while receiving substantial benefits from hosting clownfishes [28, 29], can be found solitarily [25].

The ability of clownfishes to live within the otherwise lethal tentacles of sea anemones stems from their mucus coating. The mucus coating is maintained through regular physical contact with anemone hosts, which is thought to prevent nematocyst discharge via “chemical camouflage,” causing the anemone to recognize the fish as self [30]. The long- and short-term maintenance of the symbiosis thus requires constant interaction, which should lead to regular microbial transfer among host and symbiont. In a laboratory setting, the microbial makeup of clownfish mucus was shown to change rapidly in the presence of an anemone host [31]. However, the microbial diversity of host anemones remains uncharacterized, and the multiple origins of symbiosis with clownfishes makes these animals particularly useful systems for understanding if macroscopic symbiosis generates microbial convergence across distantly related taxa.

Here, we use *in situ* field sampling to conduct the first comparative microbial study of clownfish-hosting sea anemones to test the importance of host identity, clownfish symbiont association, and habitat on the genetic and predicted functional diversity of the microbiome. We examine the microbiomes of five species of clownfish-hosting sea anemones that co-occur on a fine scale on coral reefs in the Maldives, but that vary in habitat and clownfish symbiont associations. Our five focal taxa come from three anemone clades that have evolved symbiosis with clownfishes independently [24] and therefore offer a unique opportunity to explore the interplay between evolutionary history, environment, and macroscopic symbioses in shaping the diversity of the microbiome.

## Materials and Methods

### Sample collection

Microbial samples were collected from five species of clownfish-hosting sea anemones in Huvadhoo Atoll, Republic of the Maldives (0°11’45.89”N, 73°11’3.53”E)-*Cryptodendrum adhaesivum, Entacmaea quadricolor, Heteractis aurora, H. magnifica*, and *Stichodactyla mertensii* (Table 1; Fig. S1). Focal anemones come from three clades that have evolved symbiosis with clownfishes independently: 1) Stichodactylina (*C. adhaesivum, H. magnifica*, and *S. mertensii*), 2) Heteractina (*H. aurora*), and 3) *E. quadricolor* [24]. Samples were collected from three atoll reef habitats: 1) outer atoll fore reef (10-25 m depth), 2) lagoonal fringing reef slope (5-25 m depth), and 3) shallow (1m depth) reef flat (Fig. S2). Sample localities were separated by no more than 10 km (Fig. S2). Only specimens of *H. magnifica* were present across all three habitats (Table 1). We further collected samples from two distinct *H. magnifica* phenotypes: 1) a purple phenotype found on the outer atoll fore reef and on lagoonal fringing reef slopes that hosted the Maldivian endemic clownfish *Amphiprion negripes* (Fig. S1E), and 2) a pale column phenotype found on a shallow (1 m depth) reef flat that did not host fish symbionts (Fig. S1F). All other anemone species and individuals sampled in this study hosted *Amphiprion clarkii* (Figure S1A-D), and no anemones were found without clownfishes in water beyond 1.5 m depth. Samples were collected by clipping two tentacles per individual anemone. A total of 94 tentacles from 47 individual anemones were sampled and preserved immediately in RNAlater.

**Table 1.** Clownfish-hosting sea anemones species, habitat, sample sizes, and clownfish symbiont association sampled from the Maldives and included in this study

| Species | Habitat | Sample Size<br>N = individuals<br>(tissues sequenced) | Fish symbiont |
| --- | --- | --- | --- |
| *Cryptodendrum adhaesivum* | Lagoon | N = 1 (2) | *Amphiprion clarkii* |
| *Entacmaea quadricolor* | Lagoon | N = 8 (16) | *Amphiprion clarkii* |
| *Heteractis aurora* | Fore Reef | N = 2 (4) | *Amphiprion clarkii* |
|  | Lagoon | N = 5 (10) | *Amphiprion clarkii* |
| *Heteractis magnifica* | Fore Reef | N = 7 (14) | *Amphiprion negripes* |
|  | Lagoon | N = 7 (14) | *Amphiprion negripes* |
|  | Reef Flat | N = 12 (24) | None |
| *Stichodactyla mertensii* | Fore Reef | N = 3 (6) | *Amphiprion clarkii* |
|  | Lagoon | N = 3 (6) | *Amphiprion clarkii* |
| **Total** |  | **N = 48 (96)** |  |

### DNA extraction and 16s amplicon sequencing

Genomic DNA was extracted from both tentacle samples collected from each individual using Qiagen DNeasy Blood and Tissue Kits. DNA was quantified and standardized to 5ng/uL for each sample. Microbiome diversity was assessed via Illumina sequencing targeting a 252 base pair sequence of the hypervariable v4 region of the 16S rRNA SSU gene. 16S amplicon libraries were prepared following the Earth Microbiome Protocol [32. Sequencing of 16s amplicons was conducted on an Illumina MiSeq using a V2 500 cycle kit (250 × 250 base pairs) at the National Museum of Natural History’s Laboratories of Analytical Biology.

### Microbiome data analyses

Amplicon sequence data were demultiplexed, denoised to identify amplicon sequence variants (ASVs), assembled, and analyzed using QIIME2 and associated plug-ins (see electronic supplementary material) [33]. Following demultiplexing and read-joining, we pooled reads from replicate samples to increase the number of sequence reads per individual anemone (i.e. reads from 94 tentacle samples pooled into 47 individuals). Microbial taxonomy was assigned using a Naïve Bayes classifier trained on the SILVA 132 99% database (silva-132-99-nb-classifer). Resulting feature tables were then filtered to remove ASVs that could not be identified as bacterial, and taxonomy was visualized using the QIIME2 taxa bar plot command.

To test for variation in microbial genetic diversity across host, habitat, and clownfish association, amplicon data were standardized using a rarefaction sequencing depth of 55,377 sequence reads per individual. Using QIIME2, we tested for significant (p < 0.05) variation in microbial alpha and beta genetic diversity in three sample metadata categories: 1) anemone host species, 2) clownfish symbiont association (*A. clarkii, A. negripes*, or none), and 3) habitat (atoll fore reef, reef flat, or lagoonal patch reef). Alpha diversity was calculated using Shannon’s Diversity Index (H) and *post hoc* comparisons made using non-parametric Kruskal-Wallis tests. Significant variation in beta diversity between sample categories was tested for using Bray Curtis distance measures of community dissimilarity, and *post hoc* comparisons made using permutational multivariate analysis of variance (perMANOVA). Ordination plots for beta diversity analyses were visualized using non-metric multidimensional scaling plots (nMDS).

Finally, we predicted and compared the functional diversity of the microbial metagenomes using PICRUSt (Phylogenetic Investigation of Communities by Reconstruction of Unobserved States) [34]. All ASVs were normalized for 16s copy number, and function was predicted by assigning ASVs to Kyoto Encyclopedia of Genes and Genomes Orthology categories (KEGG) [35]. Significant variation in predicted alpha and beta metagenomic functional diversity across sample metadata categories were tested for statistically using Shannon Diversity (H) and Bray-Curtis measures as above. We then used DESeq2 [36] and log-transformations to detect differentially abundant and highly variable functional groups across sample metadata categories.

## Results and Discussion

Our final dataset consisted of 47 individual anemones, > 4,500,000 sequence reads, and 6,288 ASVs. Variation in sequence reads per anemone ranged from 55,377 – 161,761 reads, with a median of 87,320 reads, and final ASV counts after rarefaction ranged from 75-797 (Fig. S3). The taxonomic composition of microbial communities varied by anemone species, clownfish symbiont association, and habitat (Fig. 1), but were largely dominated by unclassified Bacteria, Gammaproteobacteria, and to a lesser extent Bacteroidetes (Fig. 1A). Interestingly, more ASVs remained unclassified in pale column *H. magnifica* ecomorphs from reef flat habitats that did not host clownfish than for any other anemone species (Fig. 1A).

**Figure 1.**
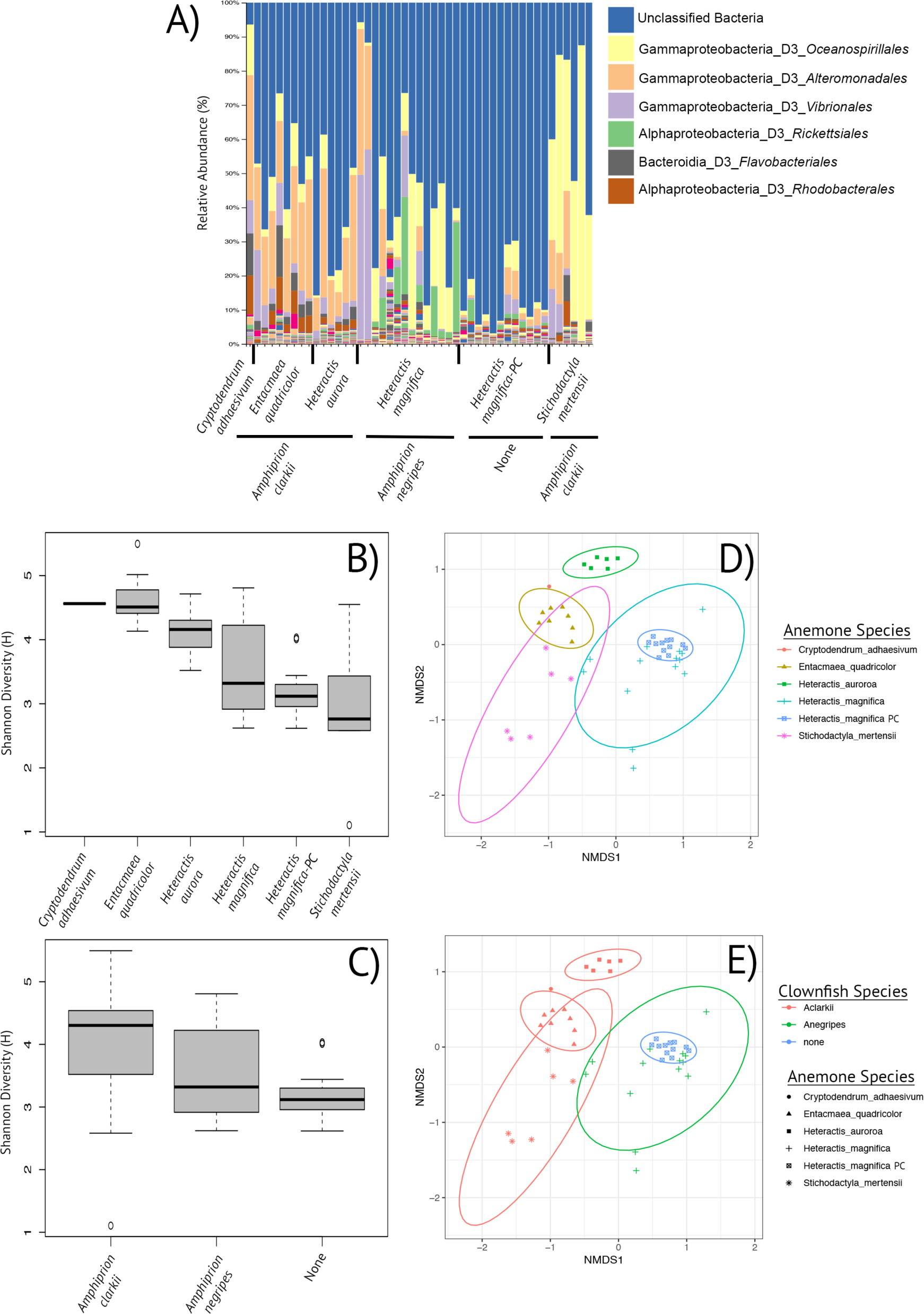
Taxonomic and genetic diversity visualizations of microbial communities from clownfish-hosting sea anemones. A) Taxonomic composition of clownfish-hosting sea anemone microbiota showing the relative abundance (%) of major microbial taxonomic groups by anemone host and clownfish symbiont association (black bars). B) Boxplot representing microbial genetic diversity (Shannon Diversity Index *H*) of the clownfish-hosting sea anemones grouped by host species. C) Boxplot representing microbial genetic diversity (Shannon Diversity Index *H*) of the clownfish-hosting sea anemones grouped by clownfish symbiont association. D) Non-metric multi-dimensional scaling (nMDS) plot of Bray-Curtis dissimilarities colored by anemone host species with 95% confidence ellipses around group centroid. E) Non-metric multi-dimensional scaling (nMDS) plot of Bray-Curtis dissimilarities of host anemone species, colored by clownfish symbiont association, with 95% confidence ellipses drawn by anemone host identity.

Alpha and beta diversity analyses indicate that anemone species, rather than clownfish symbiont association or habitat, drives the majority of the microbial genetic diversity signal in our dataset, as many of the sampled clownfish-hosting anemones harbored unique microbiomes that significantly differed from each other (Fig. 1B-E; Tables S1-6; Figs. S4 & S5). However, Bray-Curtis beta diversity analyses also indicate that anemones that host clownfish symbionts (*A. clarkii* or *A. negripes*), regardless of host or habitat, are more similar to other anemones that host the same clownfish species than they are to anemones that host a different clownfish, or that do not host fish (perMANOVA F = 15.05, p<0.002; Fig 1E; Table S5). This implies that while anemone host species seems to be primarily responsible for the microbial genetic diversity and composition, there is some degree of microbiome convergence across distantly related anemones that host the same clownfish species. The transfer of microbiota among macroscopic symbiotic partners is well demonstrated in plant-pollinator symbioses [e.g. 11, 37] and microbiome convergence has also been noted in some, but not all, leaf-cutter ant symbioses [e.g. 38-39]. Microbiome convergence among symbiotic partners is less understood in marine systems, but Pratte et al. [31] recently demonstrated changes in microbiome composition between anemone-hosting and non-hosting *A. clarkii* clownfish in a laboratory setting, implying either direct microbial transfer from anemone to clownfish or a shift in microbial diversity in response to the interaction. Our data also suggests either direct microbial transfer from the clownfish to the host, or a shift in diversity may be occurring. Pratte et al. [31] did not sample anemone microbiomes throughout the duration of their experimental trials, and we did not sample clownfish here, so it remains to be seen how convergent the microbiomes from both symbiotic partners ultimately become when in association with each other, and whether microbial transfer does occur between partners.

Functionally, alpha and beta diversity analyses using PICRUSt predicted metagenomes reinforce the role of anemone host identity in shaping the functional diversity of the host microbiome (Shannon Diversity Index, H = 28.38, p<0.0001; perMANOVA F = 14.82, p<0.002; Tables S7-8), but interestingly, these analyses also highlight that hosting clownfish symbionts increases the functional alpha and beta microbial diversity of host anemones over those that do not (H = 15.67, p<0.0001; F = 14.86, p<0.002; Fig. 2A-C; Tables S9-10). Anemones that hosted *A. clarkii* and *A. negripes* did not differ functionally from each other in either alpha or beta diversity measures (Fig. 2A-C; Table S8), and if increased functional microbial diversity is indicative of increased health as in many other taxa, these data may represent the first evidence of a previously unknown mutualistic benefit of hosting clownfishes gained by host anemones within this iconic symbiosis.

**Figure 2.**
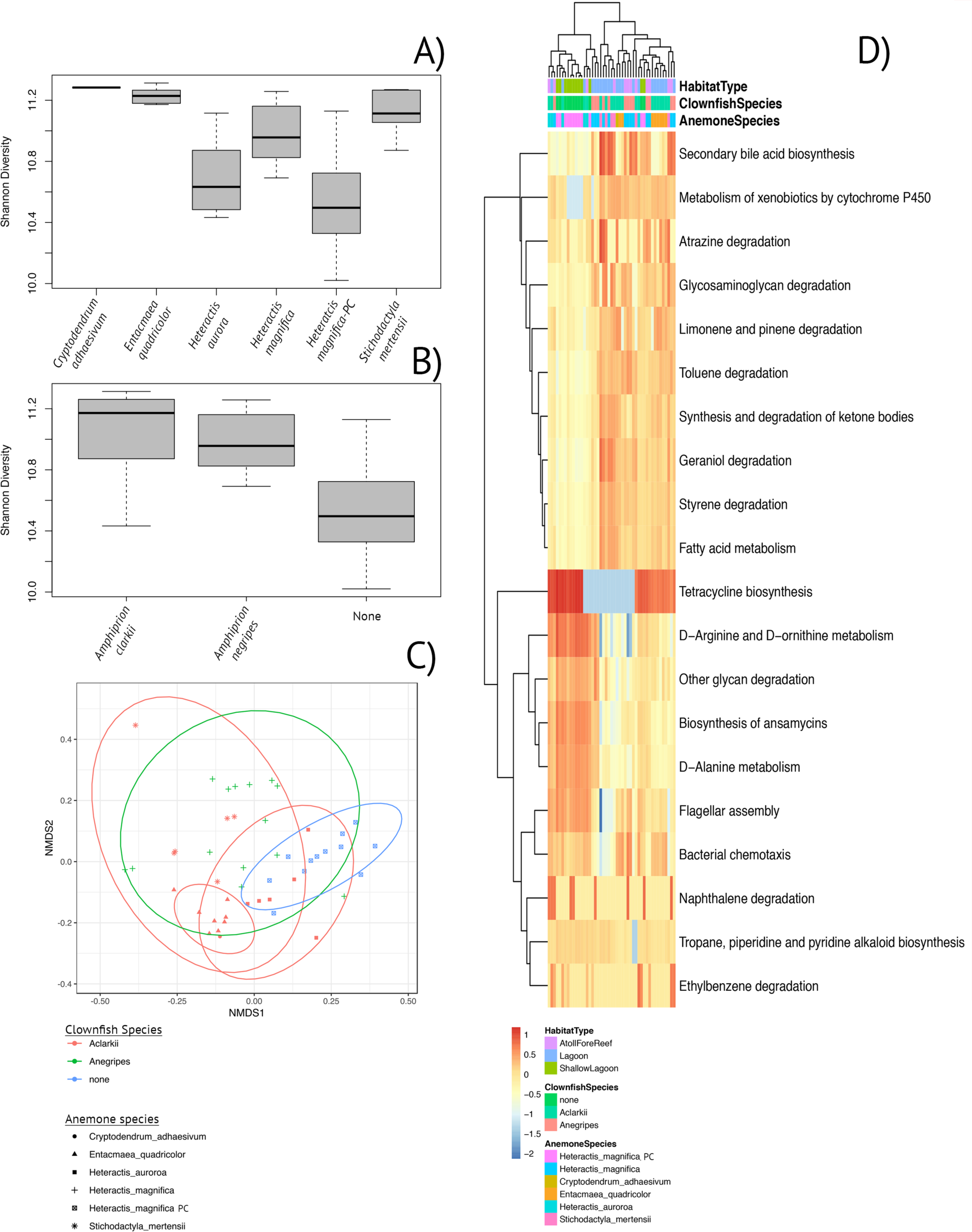
Variation in PICRUSt predicted functional microbial diversity in the clownfish hosting sea anemones. A) Boxplot representing predicted functional alpha diversity (Shannon Diversity Index *H*) of the clownfish-hosting sea anemones grouped by host species. C) Boxplot representing predicted functional alpha diversity (Shannon Diversity Index *H*) of the clownfish-hosting sea anemones grouped by clownfish symbiont association. C) Non-metric multi-dimensional scaling (nMDS) plot of predicted functional Bray-Curtis dissimilarities of host anemone species, colored by clownfish symbiont association, with 95% confidence ellipses drawn by anemone host identity. D) Heat map of top 20 most variable predicted KEGG functional categories across anemone host identity, clownfish symbiont association, and habitat.

Using DESeq2 analyses, we identified KEGG functions that were differentially abundant and highly variable between clownfish-hosting and non-hosting anemones (Table 2; Fig 2D). In clownfish-hosting anemones, Arachidonic acid (ARA) metabolic functions, part of the broader lipid metabolism pathway, were 25-fold more abundant over non-hosting anemones (Table 2). No other functional pathway was over 3-fold more abundant (Table 2, Table S11). ARA is an essential polyunsaturated fatty acid (PUFA) that could be acquired by host anemones via translocated photosynthate from their Symbiodiniaceae community, heterotrophic prey capture, or via waste byproducts from clownfish symbionts [e.g. 40-43]. Cytochrome P450 monooxygenase pathways, which catalyze ARA and other PUFAs to biologically active, intercellular signaling molecules (eicosanoids), were also highly variable and enriched primarily in host anemones (Fig. 2D). Eicosanoid lipids participate in an oxidative stress response and are hypothesized to play a role in the oxidative stress response in symbiotic cnidarians [43-44]. It is well documented that clownfish symbionts increase gas transfer and oxygenate their host anemones while also passing organic wastes to their anemone hosts, which functionally act as fertilizers for endosymbiotic Symbiodiniaceae [28-29]. It is not inconceivable that the increase in oxygen free radicals produced by fish movements through the host, and by the oxygen produced as waste during photosynthesis, could stimulate a metabolic response by the host anemone that mimics an oxidative stress response. Consequently, anemones that host fish could see a corresponding shift in microbiome diversity and function to compensate for increased ARA metabolites that could be harmful to the host. If so, these data could indicate a hidden cost of hosting mutualistic clownfishes for the anemones. However, ARA are used in a myriad of physiological processes, and its derivatives such as platelet activating factors, are also known to be involved in tissue growth and coral competition [e.g. 45-46]. More detailed microbial and metabolomic studies are needed to pinpoint the source of any increased levels of ARA in the host anemones in the presence of clownfish. Regardless of its origin here, and whether or not it reflects a beneficial aspect of the symbiosis, the degree to which microbial communities differ in ARA functions is a striking metabolic signal that the microbial communities on host sea anemones are responding to clownfish presence. Minimally, this finding is consistent with the literature that clownfish presence has a significant impact on the metabolism and physiology of the host anemone [28-29; 47-48].

**Table 2.** Top ten significantly enriched (*p*-adj <0.05) microbial KEGG Functions for clownfish hosting (gray highlighted) and non-clownfish host sea anemones. Positive log-2-fold-change values indicated microbial functions enriched in host anemones versus non-host anemones, and negative log-2-fold-change values indicated microbial functions enriched in non-host anemones versus host anemones.

| KEGG Function | log2FoldChange | $p\text{-adj}$ | Clownfish Association |
| --- | --- | --- | --- |
| Arachidonic acid metabolism | 25.841 | $2.57\text{e}^{-12}$ | Host |
| Protein digestion and absorption | 2.100 | $9.71\text{e}^{-10}$ | Host |
| Renin-angiotensin system | 1.903 | 0.040 | Host |
| Non-homologous end-joining | 1.730 | 0.007 | Host |
| Lysosome | 1.543 | 0.002 | Host |
| Phosphonate and phosphinate metabolism | 1.534 | $1.19\text{e}^{-07}$ | Host |
| Linoleic acid metabolism | 1.527 | $1.96\text{e}^{-09}$ | Host |
| Flavonoid biosynthesis | 1.516 | 0.006 | Host |
| Penicillin and cephalosporin biosynthesis | 1.431 | $9.71\text{e}^{-10}$ | Host |
| Primary bile acid biosynthesis | 1.422 | 0.001 | Host |
| RNA polymerase | -0.608 | $1.19\text{e}^{-06}$ | Non-host |
| Flagellar assembly | -0.645 | 0.005 | Non-host |
| D-Alanine metabolism | -0.711 | $2.00\text{e}^{-06}$ | Non-host |
| Biosynthesis of ansamycins | -0.740 | $5.29\text{e}^{-06}$ | Non-host |
| Insulin signaling pathway | -0.803 | $2.95\text{e}^{-06}$ | Non-host |
| D-Arginine and D-ornithine metabolism | -1.029 | 0.0008 | Non-host |
| Amoebiasis | -1.355 | 0.0003 | Non-host |
| Proteasome | -1.431 | 0.0178 | Non-host |
| Systemic lupus erythematosus | -1.798 | 0.0107 | Non-host |
| Cell adhesion molecules (CAMs) | -2.909 | 0.0017 | Non-host |

Other functional microbial groups that we recovered to be differentially abundant between host and non-host anemones provide further indication that microbial communities are responding to clownfish presence (Table 2). Bacteria involved in the renin-angiotensin system and primary bile acid biosynthesis were also more abundant in host anemones and should only be coming from, or responding to, metabolic byproducts from vertebrate sources-most likely clownfish wastes. Linoleic acid metabolic bacterial functions may also indicate an increase in the degree to which endosymbiotic Symbiodiniaceae communities in host anemones are passing PUFA photosynthates to the host-possibly reflecting a healthier endosymbiont community continually being nourished by organic clownfish waste.

In conclusion, we demonstrate for the first time that clownfish presence increases the functional diversity of the anemone host microbiome, revealing an even deeper metabolic coupling between clownfishes, host anemones, endosymbiotic Symbiodiniaceae, and what is likely a previously unrecognized mutualistic benefit of the symbiosis at the microbial level.

## Acknowledgements

We thank the Small Island Research Center (Fares-Maathoda, Maldives) for field research support and logistics. Aaron Hartmann, Melissa Ingala, Jennifer Matthews, and Susan Perkins provided valuable advice on data analysis and anthozoan metabolic functions. The Laboratories of Analytical Biology (L.A.B.) at the National Museum of Natural History for molecular services. Samples were collected under research permit 30-D/Indiv/2018/27. This work was supported by the Gerstner Scholars Postdoctoral Fellowship and the Gerstner Family Foundation, the Lerner-Gray Fund for Marine Research, and the Richard Guilder Graduate School, American Museum of Natural History to BMT, and the National Museum of Natural History to CPM.

## References

1. Boucher, D.H., ed. (1985) The Biology of Mutualism: Ecology and Evolution, Oxford University Press.

2. Margulis, L. and Fester, R., eds (1991) Symbiosis as a Source of Evolutionary Innovation, MIT Press

3. Herre, E. A., Knowlton, N., Mueller, U. G., & Rehner, S. A. (1999). The evolution of mutualisms: exploring the paths between conflict and cooperation. Trends in ecology & evolution, 14(2), 49–53.

4. Vandenkoornhuyse, P., Quaiser, A., Duhamel, M., Le Van, A., & Dufresne, A. (2015). The importance of the microbiome of the plant holobiont. New Phytologist, 206(4), 1196–1206.

5. Sweet, M. J., & Bulling, M. T. (2017). On the importance of the microbiome and pathobiome in coral health and disease. Frontiers in Marine Science, 4, 9.

6. Douglas, A. E. (1998). Nutritional interactions in insect-microbial symbioses: aphids and their symbiotic bacteria Buchnera. Annual review of entomology, 43(1), 17–37.

7. Christian, N., Whitaker, B. K., & Clay, K. (2015). Microbiomes: unifying animal and plant systems through the lens of community ecology theory. Frontiers in microbiology, 6, 869.

8. Gilbert, J.A., Quinn, R.A., Debelius, J., Xu, Z.Z., Morton, J., Garg, N., Jansson, J.K., Dorrestein, P.C. and Knight, R., 2016. Microbiome-wide association studies link dynamic microbial consortia to disease. Nature, 535(7610), p.94.

9. Barea, J. M., Pozo, M. J., Azcon, R., & Azcon-Aguilar, C. (2005). Microbial co-operation in the rhizosphere. Journal of experimental botany, 56(417), 1761–1778.

10. McFall-Ngai, M. J. (2014). The importance of microbes in animal development: lessons from the squid-vibrio symbiosis. Annual Review of Microbiology, 68, 177–194.

11. Ushio, M., Yamasaki, E., Takasu, H., Nagano, A.J., Fujinaga, S., Honjo, M.N., Ikemoto, M., Sakai, S. and Kudoh, H., 2015. Microbial communities on flower surfaces act as signatures of pollinator visitation. Scientific reports, 5, p.8695.

12. Glasl, B., Herndl, G. J., & Frade, P. R. (2016). The microbiome of coral surface mucus has a key role in mediating holobiont health and survival upon disturbance. The ISME journal, 10(9), 2280.

13. Blekhman, R., Goodrich, J.K., Huang, K., Sun, Q., Bukowski, R., Bell, J.T., Spector, T.D., Keinan, A., Ley, R.E., Gevers, D. and Clark, A.G., 2015. Host genetic variation impacts microbiome composition across human body sites. Genome biology, 16(1), p.191.

14. Smillie, C. S., Smith, M. B., Friedman, J., Cordero, O. X., David, L. A., & Alm, E. J. (2011). Ecology drives a global network of gene exchange connecting the human microbiome. Nature, 480(7376), 241.

15. Brooks, A. W., Kohl, K. D., Brucker, R. M., van Opstal, E. J., & Bordenstein, S. R. (2016). Phylosymbiosis: relationships and functional effects of microbial communities across host evolutionary history. PLoS biology, 14(11), e2000225.

16. Fonseca-García, C., Coleman-Derr, D., Garrido, E., Visel, A., Tringe, S. G., & Partida-Martínez, L. P. (2016). The cacti microbiome: interplay between habitat-filtering and host-specificity. Frontiers in microbiology, 7, 150.

17. Ainsworth, T.D., Krause, L., Bridge, T., Torda, G., Raina, J.B., Zakrzewski, M., Gates, R.D., Padilla-Gamiño, J.L., Spalding, H.L., Smith, C. and Woolsey, E.S., 2015. The coral core microbiome identifies rare bacterial taxa as ubiquitous endosymbionts. The ISME journal, 9(10), p.2261.

18. Bronstein, J. L., Alarcón, R., & Geber, M. (2006). The evolution of plant–insect mutualisms. New Phytologist, 172(3), 412–428.

19. Horká, I., De Grave, S., Fransen, C. H., Petrusek, A., & Ďuriš, Z. (2018). Multiple origins and strong phenotypic convergence in fish-cleaning palaemonid shrimp lineages. Molecular Phylogenetics & Evolution, 124, 71–81.

20. Titus, B. M., Daly, M., Vondriska, C., Hamilton, I., & Exton, D. A. (2019). Lack of strategic service provisioning by Pederson’s cleaner shrimp (Ancylomenes pedersoni) highlights independent evolution of cleaning behaviors between ocean basins. Scientific reports, 9(1), 629.

21. Litsios, G., Sims, C. A., Wüest, R. O., Pearman, P. B., Zimmermann, N. E., & Salamin, N. (2012). Mutualism with sea anemones triggered the adaptive radiation of clownfishes. BMC Evolutionary Biology, 12(1), 212.

22. Litsios, G., Pearman, P. B., Lanterbecq, D., Tolou, N., & Salamin, N. (2014). The radiation of the clownfishes has two geographical replicates. Journal of biogeography, 41(11), 2140–2149.

23. Litsios, G., & Salamin, N. (2014). Hybridisation and diversification in the adaptive radiation of clownfishes. BMC Evolutionary Biology, 14(1), 245.

24. Titus, BM, Benedict, C.*, Laroche, R.* Gusmao, LC, Van Deusen, V.,* Chiodo, T., Meyer, CP, Berumen, ML, Bartholomew, A., Yanagi, K., Reimer, JD, Fujii, T., Daly, M., Rodriguez, E. (2019) Phylogenetic relationships of the clownfish-hosting sea anemones. Molecular Phylogenetics & Evolution doi:10.1016/j.ympev.2019.106526

25. Fautin, D. G., Allen, G. R., Allen, G. R., Naturalist, A., Allen, G. R., & Naturaliste, A. (1992). Field guide to anemonefishes and their host sea anemones.

26. Huebner, L. K., Dailey, B., Titus, B. M., Khalaf, M., & Chadwick, N. E. (2012). Host preference and habitat segregation among Red Sea anemonefish: effects of sea anemone traits and fish life stages. Marine Ecology Progress Series, 464, 1–15.

27. Camp, E. F., Hobbs, J. P. A., De Brauwer, M., Dumbrell, A. J., & Smith, D. J. (2016). Cohabitation promotes high diversity of clownfishes in the Coral Triangle. Proceedings of the Royal Society B: Biological Sciences, 283(1827), 20160277.

28. Szczebak, J. T., Henry, R. P., Al-Horani, F. A., & Chadwick, N. E. (2013). Anemonefish oxygenate their anemone hosts at night. Journal of Experimental Biology, 216(6), 970–976.

29. Roopin, M., Henry, R. P., & Chadwick, N. E. (2008). Nutrient transfer in a marine mutualism: patterns of ammonia excretion by anemonefish and uptake by giant sea anemones. Marine Biology, 154(3), 547–556.

30. Mebs, D. (2009). Chemical biology of the mutualistic relationships of sea anemones with fish and crustaceans. Toxicon, 54(8), 1071–1074.

31. Pratte, Z. A., Patin, N. V., McWhirt, M. E., Caughman, A. M., Parris, D. J., & Stewart, F. J. (2018). Association with a sea anemone alters the skin microbiome of clownfish. Coral Reefs, 37(4), 1119–1125.

32. Thompson, L. R., Sanders, J. G., McDonald, D., Amir, A., Ladau, J., Locey, K. J. et al. (2017). A communal catalogue reveals Earth’s multiscale microbial diversity. Nature, 551(7681), 457.

33. Bolyen, E., Rideout, J.R., Dillon, M.R., Bokulich, N.A., Abnet, C., Al-Ghalith, G.A., Alexander, H., Alm, E.J., Arumugam, M., Asnicar, F. and Bai, Y. (2018). QIIME 2: Reproducible, interactive, scalable, and extensible microbiome data science(No. e27295v1). PeerJ Preprints.

34. Langille, M. G., Zaneveld, J., Caporaso, J. G., McDonald, D., Knights, D., Reyes, J. A., … & Beiko, R. G. (2013). Predictive functional profiling of microbial communities using 16S rRNA marker gene sequences. Nature biotechnology, 31(9), 814.

35. Kanehisa, M., & Goto, S. (2000). KEGG: kyoto encyclopedia of genes and genomes. Nucleic acids research, 28(1), 27–30.

36. Love, M. I., Huber, W., & Anders, S. (2014). Moderated estimation of fold change and dispersion for RNA-seq data with DESeq2. Genome biology, 15(12), 550.

37. McFrederick, Q. S., Thomas, J. M., Neff, J. L., Vuong, H. Q., Russell, K. A., Hale, A. R., & Mueller, U. G. (2017). Flowers and wild megachilid bees share microbes. Microbial ecology, 73(1), 188–200.

38. Aylward, F.O., Burnum, K.E., Scott, J.J., Suen, G., Tringe, S.G., Adams, S.M., Barry, K.W., Nicora, C.D., Piehowski, P.D., Purvine, S.O. and Starrett, G.J., 2012. Metagenomic and metaproteomic insights into bacterial communities in leaf-cutter ant fungus gardens. The ISME journal, 6(9), p.1688.

39. Kellner, K., Ishak, H.D., Linksvayer, T.A. and Mueller, U.G., 2015. Bacterial community composition and diversity in an ancestral ant fungus symbiosis. FEMS microbiology ecology, 91(7), p.fiv073.

40. Matthews, J.L., Oakley, C.A., Lutz, A., Hillyer, K.E., Roessner, U., Grossman, A.R., Weis, V.M. and Davy, S.K., 2018. Partner switching and metabolic flux in a model cnidarian–dinoflagellate symbiosis. Proceedings of the Royal Society B, 285(1892), p.20182336.

41. Al-Moghrabi, S., Allemand, D., Couret, J.M. and Jaubert, J., 1995. Fatty acids of the scleractinian coral Galaxea fascicularis: effect of light and feeding. Journal of Comparative Physiology B, 165(3), pp.183–192.

42. Jiang, P.L., Pasaribu, B. and Chen, C.S., 2014. Nitrogen-deprivation elevates lipid levels in Symbiodinium spp. by lipid droplet accumulation: morphological and compositional analyses. PloS one, 9(1), p.e87416.

43. Matthews, J.L., Crowder, C.M., Oakley, C.A., Lutz, A., Roessner, U., Meyer, E., Grossman, A.R., Weis, V.M. and Davy, S.K., 2017. Optimal nutrient exchange and immune responses operate in partner specificity in the cnidarian-dinoflagellate symbiosis. Proceedings of the National Academy of Sciences, 114(50), pp.13194–13199.

44. Lõhelaid, H., Teder, T. and Samel, N., 2015. Lipoxygenase-allene oxide synthase pathway in octocoral thermal stress response. Coral Reefs, 34(1), pp.143–154.

45. Quinn, R.A., Vermeij, M.J., Hartmann, A.C., Galtier d’Auriac, I., Benler, S., Haas, A., Quistad, S.D., Lim, Y.W., Little, M., Sandin, S. and Smith, J.E., 2016. Metabolomics of reef benthic interactions reveals a bioactive lipid involved in coral defence. Proceedings of the Royal Society B: Biological Sciences, 283(1829), p.20160469.

46. Galtier d’Auriac, I., Quinn, R.A., Maughan, H., Nothias, L.F., Little, M., Kapono, C.A., Cobian, A., Reyes, B.T., Green, K., Quistad, S.D. Leray, M., Smith., J.E., Dorrenstein, P.C., Rohwer, F., Deheyn, D.D., Hartmann, A.C. 2018. Before platelets: the production of platelet-activating factor during growth and stress in a basal marine organism. Proceedings of the Royal Society B: Biological Sciences, 285(1884), p.20181307.

47. Roopin, M. and Chadwick, N.E., 2009. Benefits to host sea anemones from ammonia contributions of resident anemonefish. Journal of Experimental Marine Biology and Ecology, 370(1-2), pp.27–34.

48. Cleveland, A., Verde, E.A. and Lee, R.W., 2011. Nutritional exchange in a tropical tripartite symbiosis: direct evidence for the transfer of nutrients from anemonefish to host anemone and zooxanthellae. Marine Biology, 158(3), pp.589–602.

