## Supplemental Material for "Host identity and symbiotic association affects the genetic and functional diversity of the clownfish-hosting sea anemone microbiome"

### Microbiome data analyses

Raw amplicon sequence data for each of the 94 individual samples were demultiplexed, denoised to identify amplicon sequence variants (ASVs), trimmed, and analyzed using QIIME2 [1]. Following demultiplexing, we used DADA2 [2] to denoise, join paired-end reads, and identify ASVs. Forward and reverse sequence reads were truncated using DADA2 to 200 and 180 base pairs, respectively, due to a drop in base pair quality scores past these sequence lengths (Phred score < 30). Truncated paired end reads for each sample were joined to create 252 bp consensus sequences. Because we sequenced two tentacle samples per individual, we then grouped the 94 samples into 47 individuals, thereby increasing the number of sequence reads per anemone. Microbial taxonomy was assigned using a Naïve Bayes classifier trained on the SILVA 132 99% database (silva-132-99-nb-classifer). Resulting feature tables were then filtered to remove ASVs that could not be identified as bacterial. To visualize taxonomy, the QIIME2 taxa bar plot command was used.

Variation in microbial genetic diversity across host, habitat, and clownfish association was tested for statistically using QIIME2 plugins. To account for variation in sequence read depth across individual anemones, sequences were standardized using rarefaction curves. Our final dataset was rarified to 55,377 sequence reads per individual, in order to retain microbial data from all 47 sequenced anemones. Alpha diversity was calculated using Shannon's Diversity Index (H), which accounts for species richness and evenness within each sample category. Shannon Diversity measures were calculated and visualized using the q2-diversity plugin in QIIME2. For alpha diversity analyses we tested for variation in microbial diversity in three sample categories: 1) anemone host, 2) clownfish symbiont association, and 3) habitat. Statistical significance ($p < 0.05$) was assessed for all groups, and each pairwise comparison, using non-parametric Kruskal-Wallis tests.

Significant variation in beta diversity between sample categories was tested for using Bray Curtis distance measures of community dissimilarity, as well as unweighted UniFrac distance measures that incorporate phylogenetic relatedness between microbial taxa. Like above, we tested for variation in microbial beta diversity in three sample categories: 1) anemone host, 2) clownfish symbiont association, and 3) habitat. Statistical significance ($p < 0.05$) was assessed using permutational multivariate analysis of variance (perMANOVA) on both Bray Curtis and unweighted UniFrac distances for all three sample categories. Ordination plots for Bray Curtis and unweighted UniFrac beta diversity analyses were visualized in the context of sample metadata using Principal Coordinates (PCoA). All beta diversity analyses were conducted using the q2-diversity plugin in QIIME2. In order to perform both alpha and beta diversity measures that incorporate phylogenetic relatedness, we used the QIIME2 plugin q2-phylogeny to generate a masked sequence alignment with MAFFT [3] to remove positions that are highly variable, and a mid-point rooted phylogenetic tree using FastTree [4].

**References**

1. Bolyen, E., Rideout, J.R., Dillon, M.R., Bokulich, N.A., Abnet, C., Al-Ghalith, G.A., Alexander, H., Alm, E.J., Arumugam, M., Asnicar, F. and Bai, Y., 2018. *QIIME 2: Reproducible, interactive, scalable, and extensible microbiome data science*(No. e27295v1). PeerJ Preprints.
2. Callahan, B. J., McMurdie, P. J., Rosen, M. J., Han, A. W., Johnson, A. J. A., & Holmes, S. P. (2016). DADA2: high-resolution sample inference from Illumina amplicon data. *Nature methods*, *13*(7), 581.
3. Katoh, K., & Standley, D. M. (2013). MAFFT multiple sequence alignment software version 7: improvements in performance and usability. *Molecular biology and evolution*, *30*(4), 772-780.
4. Price, M. N., Dehal, P. S., & Arkin, A. P. (2009). FastTree: computing large minimum evolution trees with profiles instead of a distance matrix. *Molecular biology and evolution*, *26*(7), 1641-1650.

### Supplementary Tables

Table S1: Kruskal-Wallis *post-hoc* pairwise comparisons of Shannon Diversity Indices (H) among the clownfish-hosting sea anemone microbial communities based on host identity. Each comparison is listed along with sample sizes, H-index, p-value (significant values of $p<0.05$ are bolded), and q-values (Benjamini & Hochberg adjusted p-values; significant values of $q<0.05$ are bolded). Overall Kruskall-Wallis test for all groups was significant for anemone host ($H = 20.06$, $p<0.002$).

| **Group 1** | **Group 2** | **H** | **p-value** | **q-value** |
|---|---|---|---|---|
| Cryptodendrum_adhaesivum (n=1) | Entacmaea_quadricolor (n=8) | 0.60 | 0.44 | 0.47 |
| Cryptodendrum_adhaesivum (n=1) | Heteractis_auroroa (n=6) | 1.00 | 0.32 | 0.40 |
| Cryptodendrum_adhaesivum (n=1) | Heteractis_magnifica (n=14) | 0.86 | 0.35 | 0.41 |
| Cryptodendrum_adhaesivum (n=1) | Heteractis_magnifica_PC (n=12) | 2.57 | 0.11 | 0.20 |
| Cryptodendrum_adhaesivum (n=1) | Stichodactyla_mertensii (n=6) | 2.25 | 0.13 | 0.20 |
| Entacmaea_quadricolor (n=8) | Heteractis_auroroa (n=6) | 4.27 | **0.04** | 0.10 |
| Entacmaea_quadricolor (n=8) | Heteractis_magnifica (n=14) | 6.38 | **0.01** | 0.06 |
| Entacmaea_quadricolor (n=8) | Heteractis_magnifica_PC (n=12) | 13.71 | **0.00** | **0.00** |
| Entacmaea_quadricolor (n=8) | Stichodactyla_mertensii (n=6) | 5.40 | **0.02** | 0.08 |
| Heteractis_auroroa (n=6) | Heteractis_magnifica (n=14) | 2.46 | 0.12 | 0.20 |
| Heteractis_auroroa (n=6) | Heteractis_magnifica_PC (n=12) | 8.98 | **0.00** | **0.02** |
| Heteractis_auroroa (n=6) | Stichodactyla_mertensii (n=6) | 4.33 | **0.04** | 0.10 |
| Heteractis_magnifica (n=14) | Heteractis_magnifica_PC (n=12) | 0.52 | 0.47 | 0.47 |
| Heteractis_magnifica (n=14) | Stichodactyla_mertensii (n=6) | 2.72 | 0.10 | 0.20 |
| Heteractis_magnifica_PC (n=12) | Stichodactyla_mertensii (n=6) | 1.26 | 0.26 | 0.36 |

Table S2. Kruskal-Wallis *post-hoc* pairwise comparisons of Shannon Diversity Indices (H) among the clownfish-hosting sea anemone microbial communities based on clownfish symbiont association. Each comparison is listed along with sample sizes, H-index, p-value (significant values of p<0.05 are bolded), and q-values (Benjamini & Hochberg adjusted p-values; significant values of q<0.05 are bolded). Overall Kruskall-Wallis test for all groups was significant for clownfish symbiont association (H = 6.98, p<0.05).

| **Group 1** | **Group 2** | **H** | **p-value** | **q-value** |
|---|---|---|---|---|
| Amphiprion_clarkii (n=21) | Amphiprion_negripes (n=14) | 1.81 | 0.18 | 0.27 |
| Amphiprion_clarkii (n=21) | none (n=12) | 7.67 | **0.01** | **0.02** |
| Amphiprion_negripes (n=14) | none (n=12) | 0.52 | 0.47 | 0.47 |

Table S3. Kruskal-Wallis *post-hoc* pairwise comparisons of Shannon Diversity Indices (H) among the clownfish-hosting sea anemone microbial communities based on habitat. Each comparison is listed along with sample sizes, H-index, p-value (significant values of p<0.05 are bolded), and q-values (Benjamini & Hochberg adjusted p-values; significant values of q<0.05 are bolded). Overall Kruskall-Wallis test for all groups was non-significant for habitat (H = 5.02, p=0.08).

| **Group 1** | **Group 2** | **H** | **p-value** | **q-value** |
|---|---|---|---|---|
| Fore Reef (n=12) | Lagoonal Patch Reef (n=23) | 0.10 | 0.75 | 0.75 |
| Fore Reef (n=12) | Reef Flat (n=12) | 7.05 | **0.01** | **0.02** |
| Lagoon Patch Reef (n=23) | Reef Flat (n=12) | 2.13 | 0.14 | 0.22 |

Table S4. perMANOVA *post-hoc* pairwise comparisons of Bray-Curtis dissimilarities among the clownfish-hosting sea anemone microbial communities based on host identity. Each comparison is listed along with sample sizes, pseudo-F, p-value (significant values of p<0.05 are bolded), and q-values (Benjamini & Hochberg adjusted p-values; significant values of q<0.05 are bolded). perMANOVA tests were based on 999 permutations. Overall perMANOVA test for all groups was significant for host identity (F = 18.83, p<0.002).

| **Group 1** | **Group 2** | **N** | **pseudo-F** | **p-value** | **q-value** |
|---|---|---|---|---|---|
| Cryptodendrum_adhaesivum | Entacmaea_quadricolor | 9 | 4.36 | 0.10 | 0.12 |
| Cryptodendrum_adhaesivum | Heteractis_auroroa | 7 | 7.63 | 0.14 | 0.14 |
| Cryptodendrum_adhaesivum | Heteractis_magnifica | 15 | 2.55 | 0.07 | 0.09 |
| Cryptodendrum_adhaesivum | Heteractis_magnifica_PC | 13 | 59.21 | 0.06 | 0.09 |
| Cryptodendrum_adhaesivum | Stichodactyla_mertensii | 7 | 5.44 | 0.14 | 0.14 |

| | | | | | |
|---|---|---|---|---|---|
| Entacmaea_quadricolor | Heteractis_auroroa | 14 | 25.98 | **0.00** | **0.00** |
| Entacmaea_quadricolor | Heteractis_magnifica | 22 | 16.19 | **0.00** | **0.00** |
| Entacmaea_quadricolor | Heteractis_magnifica_PC | 20 | 95.08 | **0.00** | **0.00** |
| Entacmaea_quadricolor | Stichodactyla_mertensii | 14 | 23.58 | **0.00** | **0.00** |
| Heteractis_auroroa | Heteractis_magnifica | 20 | 13.81 | **0.00** | **0.00** |
| Heteractis_auroroa | Heteractis_magnifica_PC | 18 | 116.08 | **0.00** | **0.00** |
| Heteractis_auroroa | Stichodactyla_mertensii | 12 | 24.97 | **0.00** | **0.00** |
| Heteractis_magnifica | Heteractis_magnifica_PC | 26 | 6.48 | **0.00** | **0.00** |
| Heteractis_magnifica | Stichodactyla_mertensii | 20 | 12.61 | **0.00** | **0.00** |
| Heteractis_magnifica_PC | Stichodactyla_mertensii | 18 | 78.23 | **0.00** | **0.00** |

Table S5. perMANOVA *post-hoc* pairwise comparisons of Bray-Curtis dissimilarities among the clownfish-hosting sea anemone microbial communities based on clownfish symbiont association. Each comparison is listed along with sample sizes, pseudo-F, p-value (significant values of p<0.05 are bolded), and q-values (Benjamini & Hochberg adjusted p-values; significant values of q<0.05 are bolded). perMANOVA tests were based on 999 permutations. Overall perMANOVA test for all groups was significant for clownfish symbiont association (F = 15.05, p<0.002).

| **Group 1** | **Group 2** | **N** | **pseudo-F** | **p-value** | **q-value** |
|---|---|---|---|---|---|
| Amphiprion_clarkii | Amphiprion_negripes | 35 | 12.038 | **0.001** | **0.001** |
| Amphiprion_clarkii | none | 33 | 24.253 | **0.001** | **0.001** |
| Amphiprion_negripes | none | 26 | 6.482 | **0.001** | **0.001** |

Table S6. perMANOVA *post-hoc* pairwise comparisons of Bray-Curtis dissimilarities among the clownfish-hosting sea anemone microbial communities based on habitat. Each comparison is listed along with sample sizes, pseudo-F, p-value (significant values of p<0.05 are bolded), and q-values (Benjamini & Hochberg adjusted p-values; significant values of q<0.05 are bolded). perMANOVA tests were based on 999 permutations. Overall perMANOVA test for all groups was significant for habitat (F = 6.41, p<0.002).

| **Group 1** | **Group 2** | **N** | **pseudo-F** | **p-value** | **q-value** |
|---|---|---|---|---|---|
| Fore Reef | Lagoonal Patch Reef | 35 | 1.409280757 | 0.189 | 0.189 |
| Fore Reef | Reef Flat | 24 | 9.116064829 | **0.001** | **0.0015** |
| Lagoonal Patch Reef | Reef Flat | 35 | 12.35788564 | **0.001** | **0.0015** |

Table S7. Kruskal-Wallis *post-hoc* pairwise comparisons of PICRUSt predicted functional Shannon Diversity Indices (H) among the clownfish-hosting sea anemone microbial communities based on host identity. Each comparison is listed along with sample sizes, H-index, p-value (significant values of p<0.05 are bolded), and q-values (Benjamini & Hochberg adjusted p-values; significant values of q<0.05 are bolded). Overall Kruskall-Wallis test for all groups was significant for anemone host (H = 28.38, p<0.00001).

| **Group 1** | **Group 2** | **H** | **p-value** | **q-value** |
|---|---|---|---|---|
| Cryptodendrum_adhaesivum (n=1) | Entacmaea_quadricolor (n=8) | 1.35 | 0.25 | 0.26 |
| Cryptodendrum_adhaesivum (n=1) | Heteractis_auroroa (n=6) | 2.25 | 0.13 | 0.15 |
| Cryptodendrum_adhaesivum (n=1) | Heteractis_magnifica (n=14) | 2.63 | 0.11 | 0.15 |
| Cryptodendrum_adhaesivum (n=1) | Heteractis_magnifica_PC (n=12) | 2.57 | 0.11 | 0.15 |
| Cryptodendrum_adhaesivum (n=1) | Stichodactyla_mertensii (n=6) | 2.25 | 0.13 | 0.15 |
| Entacmaea_quadricolor (n=8) | Heteractis_auroroa (n=6) | 9.60 | **0.00** | **0.01** |
| Entacmaea_quadricolor (n=8) | Heteractis_magnifica (n=14) | 8.22 | **0.00** | **0.01** |
| Entacmaea_quadricolor (n=8) | Heteractis_magnifica_PC (n=12) | 13.71 | **0.00** | **0.00** |
| Entacmaea_quadricolor (n=8) | Stichodactyla_mertensii (n=6) | 2.40 | 0.12 | 0.15 |
| Heteractis_auroroa (n=6) | Heteractis_magnifica (n=14) | 4.25 | **0.04** | **0.08** |
| Heteractis_auroroa (n=6) | Heteractis_magnifica_PC (n=12) | 0.88 | 0.35 | 0.35 |
| Heteractis_auroroa (n=6) | Stichodactyla_mertensii (n=6) | 5.03 | **0.02** | 0.06 |
| Heteractis_magnifica (n=14) | Heteractis_magnifica_PC (n=12) | 10.17 | **0.00** | **0.01** |
| Heteractis_magnifica (n=14) | Stichodactyla_mertensii (n=6) | 2.46 | 0.12 | 0.15 |
| Heteractis_magnifica_PC (n=12) | Stichodactyla_mertensii (n=6) | 7.89 | **0.00** | **0.01** |

Table S8. perMANOVA *post-hoc* pairwise comparisons of PICRUSt predicted functional Bray-Curtis dissimilarities among the clownfish-hosting sea anemone microbial communities based on host identity. Each comparison is listed along with sample sizes, pseudo-F, p-value (significant values of $p<0.05$ are bolded), and q-values (Benjamini & Hochberg adjusted p-values; significant values of $q<0.05$ are bolded). perMANOVA tests were based on 999 permutations. Overall perMANOVA test for all groups was significant for host identity ($F = 14.86$, $p<0.002$).

| **Group 1** | **Group 2** | **N** | **pseudo-F** | **p-value** | **q-value** |
|---|---|---|---|---|---|
| Cryptodendrum_adhaesivum | Entacmaea_quadricolor | 9 | 1.26 | 0.25 | 0.27 |
| Cryptodendrum_adhaesivum | Heteractis_auroroa | 7 | 10.04 | 0.12 | 0.17 |
| Cryptodendrum_adhaesivum | Heteractis_magnifica | 15 | 1.58 | 0.27 | 0.27 |
| Cryptodendrum_adhaesivum | Heteractis_magnifica_PC | 13 | 13.42 | 0.08 | 0.12 |
| Cryptodendrum_adhaesivum | Stichodactyla_mertensii | 7 | 5.37 | 0.15 | 0.18 |
| Entacmaea_quadricolor | Heteractis_auroroa | 14 | 38.40 | **0.00** | **0.00** |
| Entacmaea_quadricolor | Heteractis_magnifica | 22 | 7.70 | **0.00** | **0.00** |
| Entacmaea_quadricolor | Heteractis_magnifica_PC | 20 | 65.52 | **0.00** | **0.00** |
| Entacmaea_quadricolor | Stichodactyla_mertensii | 14 | 21.63 | **0.00** | **0.00** |
| Heteractis_auroroa | Heteractis_magnifica | 20 | 7.39 | **0.00** | **0.00** |
| Heteractis_auroroa | Heteractis_magnifica_PC | 18 | 1.80 | 0.18 | 0.20 |
| Heteractis_auroroa | Stichodactyla_mertensii | 12 | 38.48 | **0.00** | **0.00** |
| Heteractis_magnifica | Heteractis_magnifica_PC | 26 | 17.39 | **0.00** | **0.00** |
| Heteractis_magnifica | Stichodactyla_mertensii | 20 | 3.56 | **0.03** | 0.06 |
| Heteractis_magnifica_PC | Stichodactyla_mertensii | 18 | 55.15 | **0.00** | **0.00** |

Table S9. Kruskal-Wallis *post-hoc* pairwise comparisons of PICRUSt predicted functional Shannon Diversity Indices (H) among the clownfish-hosting sea anemone microbial communities based on clownfish symbiont association. Each comparison is listed along with sample sizes, H-index, p-value (significant values of $p<0.05$ are bolded), and q-values (Benjamini & Hochberg adjusted p-values; significant values of $q<0.05$ are bolded). Overall Kruskall-Wallis test for all groups was significant for anemone host ($H = 15.67$, $p<0.0001$).

| **Group 1** | **Group 2** | **H** | **p-value** | **q-value** |
|---|---|---|---|---|
| Amphiprion_clarkii (n=21) | Amphiprion_negripes (n=14) | 2.10 | 0.15 | 0.15 |
| Amphiprion_clarkii (n=21) | none (n=12) | 12.38 | **0.00** | **0.00** |
| Amphiprion_negripes (n=14) | none (n=12) | 10.17 | **0.00** | **0.00** |

Table S10. perMANOVA *post-hoc* pairwise comparisons of PICRUSt predicted functional Bray-Curtis dissimilarities among the clownfish-hosting sea anemone microbial communities based on clownfish symbiont association. Each comparison is listed along with sample sizes, pseudo-F, p-value (significant values of $p<0.05$ are bolded), and q-values (Benjamini & Hochberg adjusted p-values; significant values of $q<0.05$ are bolded). perMANOVA tests were based on 999 permutations. Overall perMANOVA test for all groups was significant for host identity ($F = 11.40$, $p<0.002$).

| Group 1 | Group 2 | N | pseudo-F | p-value | q-value |
|---|---|---|---|---|---|
| Amphiprion_clarkii | Amphiprion_negripes | 35 | 1.77 | 0.16 | 0.16 |
| Amphiprion_clarkii | none | 33 | 20.41 | **0.00** | **0.00** |
| Amphiprion_negripes | none | 26 | 17.39 | **0.00** | **0.00** |

Table S11. Significantly enriched (*p*-adj <0.05) microbial KEGG Functions for clownfish hosting and non-clownfish host sea anemones. Positive log-2-fold-change values indicated microbial functions enriched in host anemones versus non-host anemones, and negative log-2-fold-change values indicated microbial functions enriched in non-host anemones versus host anemones.

| KEGG Function | log2FoldChange | *p*-adj |
|---|---|---|
| Arachidonic acid metabolism | 25.84060812 | 2.57E-12 |
| Protein digestion and absorption | 2.09997851 | 9.71E-10 |
| Renin-angiotensin system | 1.90258601 | 0.039985632 |
| Non-homologous end-joining | 1.730452832 | 0.006824289 |
| Lysosome | 1.543359537 | 0.001625682 |
| Phosphonate and phosphinate metabolism | 1.533530999 | 1.19E-07 |
| Linoleic acid metabolism | 1.526685024 | 1.96E-09 |
| Flavonoid biosynthesis | 1.515829143 | 0.00557303 |
| Penicillin and cephalosporin biosynthesis | 1.431310866 | 9.71E-10 |
| Primary bile acid biosynthesis | 1.421856866 | 0.001462482 |
| Nitrotoluene degradation | 1.407047758 | 3.05E-07 |
| Secondary bile acid biosynthesis | 1.36119333 | 0.002207116 |
| African trypanosomiasis | 1.337946027 | 0.011055476 |
| Styrene degradation | 1.312549228 | 1.15E-07 |
| Toluene degradation | 1.252998538 | 3.46E-08 |
| Steroid hormone biosynthesis | 1.225459212 | 0.000743146 |
| Apoptosis | 1.218959474 | 0.001993251 |
| Steroid biosynthesis | 1.214822146 | 0.003252725 |
| Dioxin degradation | 1.052516033 | 0.005374759 |
| Peroxisome | 0.943285201 | 1.19E-07 |

| | | |
|---|---|---|
| Chlorocyclohexane and chlorobenzene degradation | 0.935245288 | 0.027965918 |
| Geraniol degradation | 0.898767657 | 2.64E-07 |
| Fluorobenzoate degradation | 0.875715514 | 0.031765262 |
| Fatty acid metabolism | 0.799845019 | 1.48E-07 |
| Benzoate degradation | 0.761560462 | 1.38E-07 |
| Biosynthesis of siderophore group nonribosomal peptides | 0.754358266 | 0.026101142 |
| Butanoate metabolism | 0.745244067 | 2.74E-08 |
| Tryptophan metabolism | 0.724493789 | 6.17E-09 |
| Synthesis and degradation of ketone bodies | 0.643446006 | 1.94E-07 |
| Biosynthesis of unsaturated fatty acids | 0.595594623 | 1.70E-08 |
| Valine, leucine and isoleucine degradation | 0.575522626 | 7.74E-09 |
| Ubiquinone and other terpenoid-quinone biosynthesis | 0.561489978 | 5.36E-08 |
| Glutathione metabolism | 0.452920612 | 6.61E-07 |
| Chloroalkane and chloroalkene degradation | 0.418087364 | 3.72E-05 |
| Lysine degradation | 0.360947681 | 1.20E-05 |
| Drug metabolism - other enzymes | 0.34932818 | 5.10E-06 |
| ABC transporters | 0.330094305 | 0.000787972 |
| beta-Alanine metabolism | 0.320329772 | 0.000557669 |
| Taurine and hypotaurine metabolism | 0.308152166 | 2.54E-05 |
| Tyrosine metabolism | 0.271323954 | 5.33E-05 |
| Sulfur relay system | 0.255256234 | 1.19E-07 |
| Caprolactam degradation | 0.243209237 | 0.000485513 |
| Aminobenzoate degradation | 0.217087626 | 0.017072806 |
| Propanoate metabolism | 0.21511719 | 3.16E-05 |
| Nitrogen metabolism | 0.201824934 | 0.003175906 |
| Glyoxylate and dicarboxylate metabolism | 0.196538079 | 4.69E-05 |
| Ribosome biogenesis in eukaryotes | 0.138157552 | 0.010066622 |
| RNA transport | 0.13575734 | 0.033568322 |
| Nicotinate and nicotinamide metabolism | 0.096261857 | 0.000485513 |
| Base excision repair | 0.08537287 | 0.00679967 |
| Selenocompound metabolism | -0.084737251 | 0.019503792 |
| Glycolysis / Gluconeogenesis | -0.095645643 | 0.000436916 |
| Sulfur metabolism | -0.105914286 | 0.020837774 |
| Homologous recombination | -0.106396489 | 0.042642772 |
| Citrate cycle (TCA cycle) | -0.110228636 | 0.009011031 |
| Mismatch repair | -0.112063756 | 0.011055476 |
| Carbon fixation pathways in prokaryotes | -0.112229408 | 5.33E-05 |
| Pyrimidine metabolism | -0.123236725 | 0.000367458 |
| Nucleotide excision repair | -0.161026162 | 0.003252725 |

| | | |
|---|---|---|
| Cysteine and methionine metabolism | -0.175771806 | 3.81E-05 |
| Cell cycle - Caulobacter | -0.1819717 | 0.00843676 |
| RNA degradation | -0.184987572 | 2.76E-05 |
| Starch and sucrose metabolism | -0.189395204 | 0.000553123 |
| Alanine, aspartate and glutamate metabolism | -0.238532289 | 4.51E-07 |
| Terpenoid backbone biosynthesis | -0.239034378 | 8.58E-06 |
| Carbon fixation in photosynthetic organisms | -0.241747554 | 6.87E-06 |
| Peptidoglycan biosynthesis | -0.275780139 | 2.34E-05 |
| One carbon pool by folate | -0.277527506 | 2.96E-06 |
| Pentose phosphate pathway | -0.277543669 | 2.50E-05 |
| Amino sugar and nucleotide sugar metabolism | -0.287327961 | 2.96E-06 |
| Ribosome | -0.295826929 | 7.93E-05 |
| Lysine biosynthesis | -0.30845282 | 6.87E-06 |
| Thiamine metabolism | -0.319636226 | 8.48E-07 |
| Biosynthesis of vancomycin group antibiotics | -0.326152934 | 0.000838933 |
| Photosynthesis | -0.327242566 | 0.000112531 |
| Fatty acid biosynthesis | -0.332179712 | 2.64E-07 |
| Phenylalanine, tyrosine and tryptophan biosynthesis | -0.347012689 | 2.64E-07 |
| Aminoacyl-tRNA biosynthesis | -0.349099038 | 1.32E-05 |
| Zeatin biosynthesis | -0.351569252 | 3.00E-05 |
| Lipoic acid metabolism | -0.364084658 | 1.58E-05 |
| Galactose metabolism | -0.364316006 | 0.004295064 |
| Pantothenate and CoA biosynthesis | -0.38088229 | 6.07E-07 |
| Biotin metabolism | -0.38146845 | 2.02E-05 |
| Valine, leucine and isoleucine biosynthesis | -0.393860255 | 1.25E-06 |
| Lipopolysaccharide biosynthesis | -0.418620866 | 9.83E-07 |
| D-Glutamine and D-glutamate metabolism | -0.420865427 | 7.88E-07 |
| Sphingolipid metabolism | -0.471443437 | 0.001246726 |
| Streptomycin biosynthesis | -0.479713633 | 1.00E-05 |
| Fructose and mannose metabolism | -0.55124917 | 2.81E-07 |
| Other glycan degradation | -0.599223605 | 0.000924724 |
| RNA polymerase | -0.608109957 | 1.19E-06 |
| Flagellar assembly | -0.644595554 | 0.004662413 |
| D-Alanine metabolism | -0.710604584 | 2.00E-06 |
| Biosynthesis of ansamycins | -0.739601695 | 5.29E-06 |
| Insulin signaling pathway | -0.802557986 | 2.95E-06 |
| D-Arginine and D-ornithine metabolism | -1.028839821 | 0.000785585 |
| Amoebiasis | -1.3550355 | 0.000342242 |
| Proteasome | -1.430869603 | 0.017775139 |
| Systemic lupus erythematosus | -1.798203039 | 0.010713683 |

| | | |
|---|---|---|
| Cell adhesion molecules (CAMs) | -2.908698738 | 0.001741795 |

### Supplementary Figures

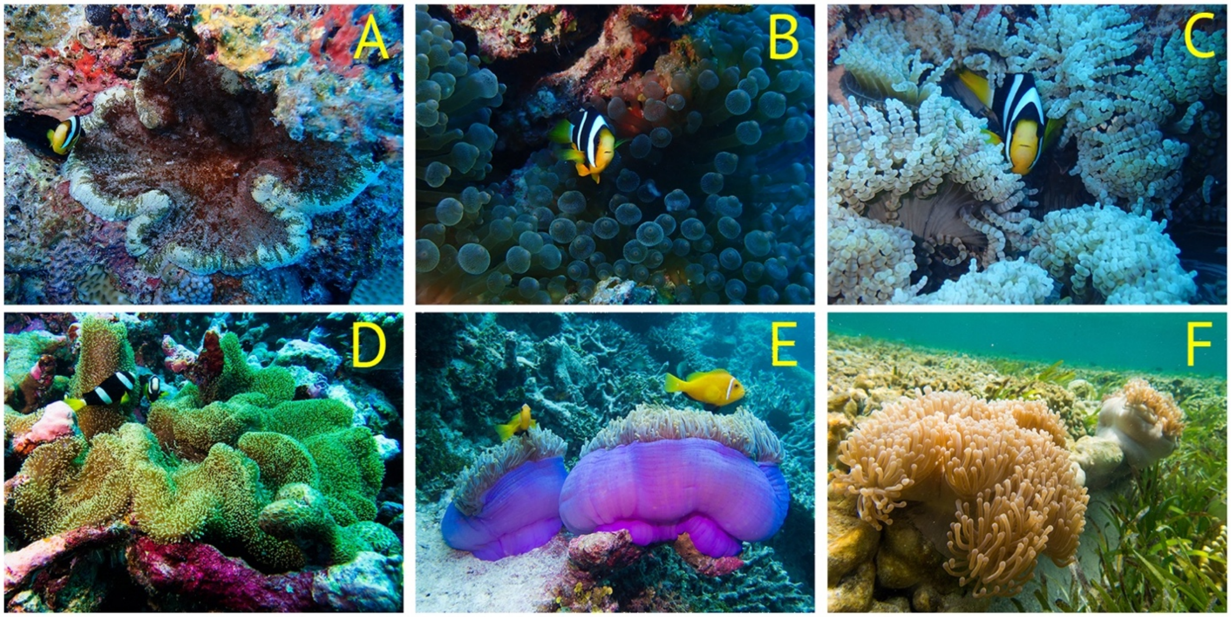


Figure S1. Representative images of the clownfish-hosting sea anemones sampled from Huvadhoo Atoll, Republic of the Maldives. A) *Cryptodendrum adhaesivum*, B) *Entacmaea quadricolor*, C) *Heteractis aurora*, D) *Stichodactyla mertensii*, E) *Heteractis magnifica*- reef slope phenotype with fish, F) *Heteractis magnifica*- Pale Column (PC) reef flat phenotype with no fish. Clownfish symbionts in panels A-D are *Amphiprion clarkii*, and in panel E is *Amphiprion negripes*.

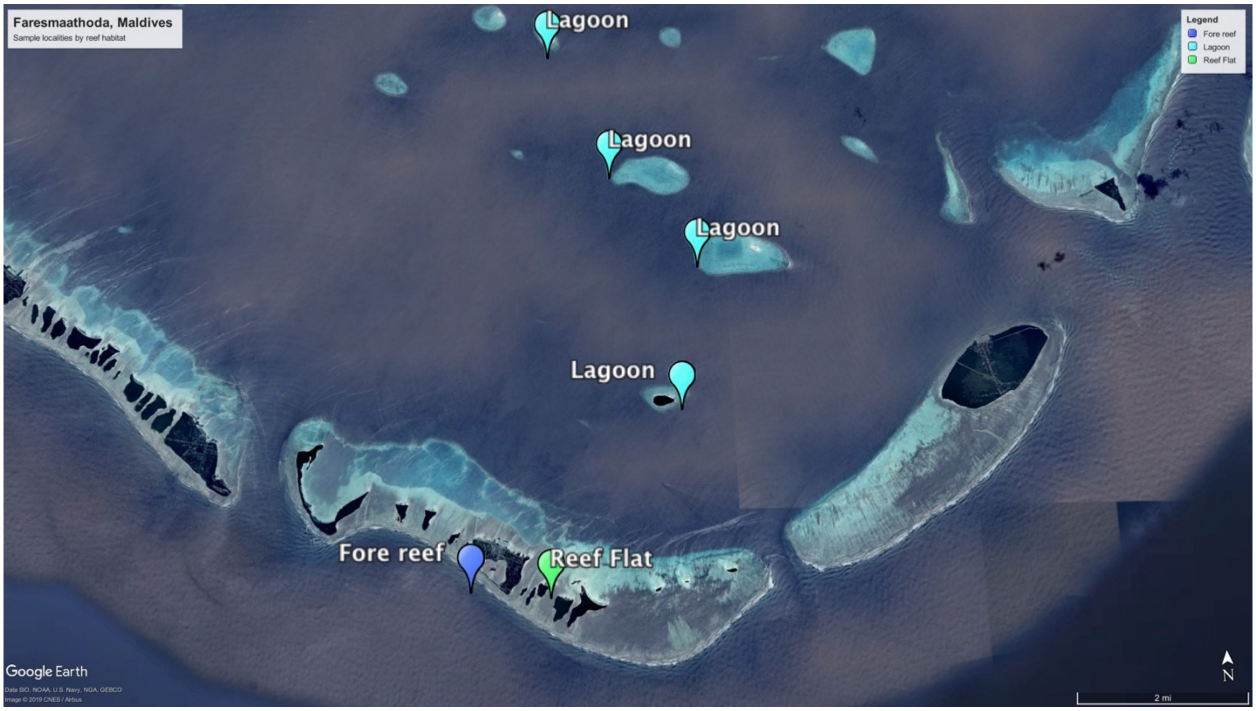


Figure S2. Satellite image of sample localities in Huvadhoo Atoll, Republic of the Maldives. Sample localities are labeled by broad reef habitat: Royal Blue = Fore Reef, Green = Reef Flat, and Light Blue = Lagoon.

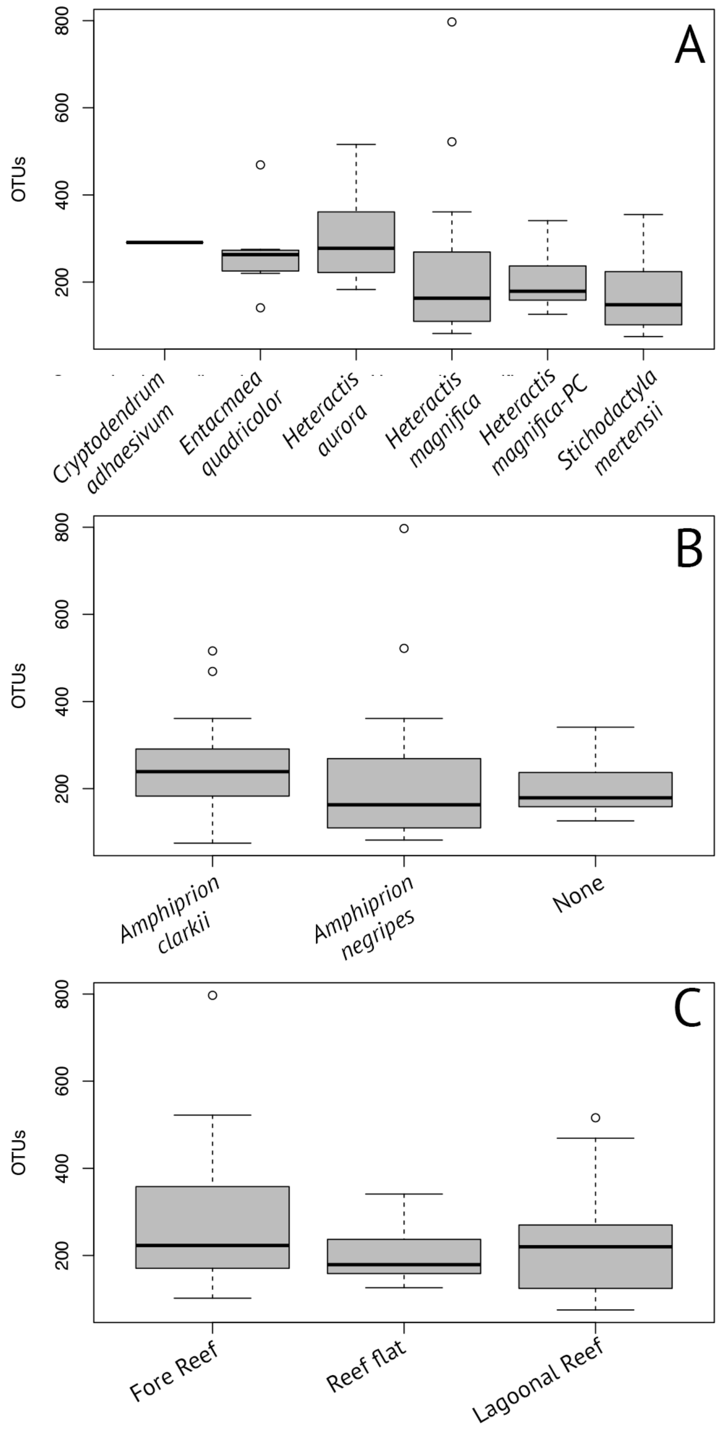


Figure S3. Distribution of microbial Amplicon Sequence Variant counts after sample rarefaction by A) anemone host species, B) clownfish symbiont association, and C) habitat. Individuals were standardized using a rarefaction sequencing depth of 55,377.

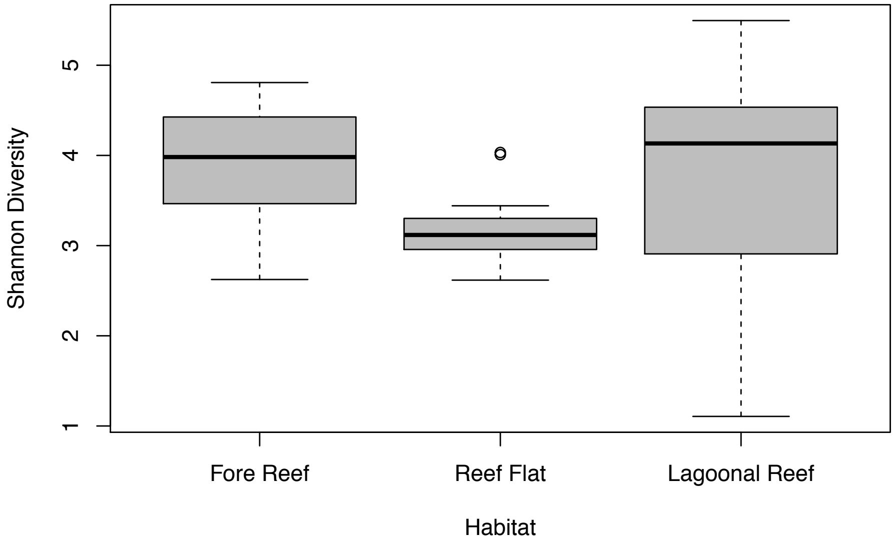


Figure S4. Boxplot representing microbial genetic diversity (Shannon Diversity Index *H*) of the clownfish-hosting sea anemones grouped by habitat.

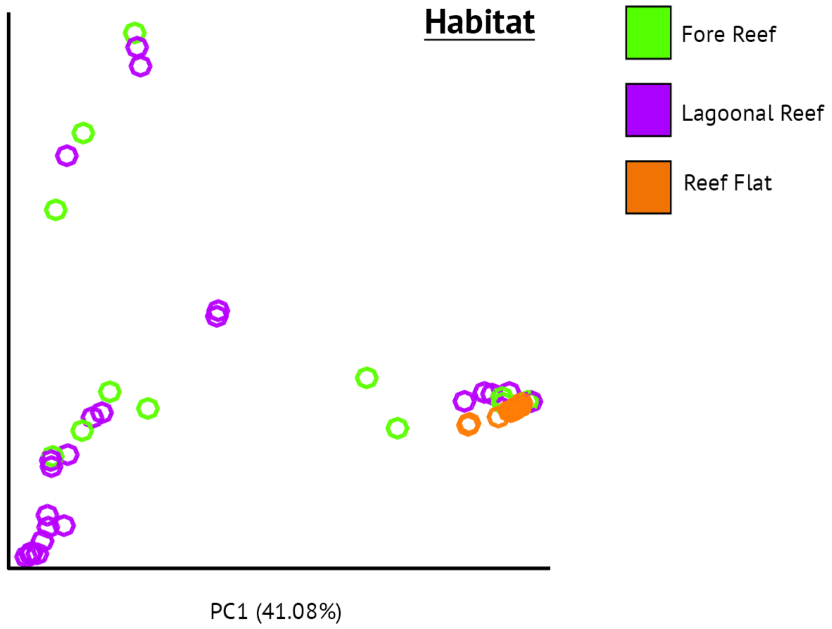


Figure S5. Non-metric multi-dimensional scaling (nMDS) plot of Bray-Curtis dissimilarities colored by coral reef habitat.
